# BoundaryStats: An R package to calculate boundary overlap statistics

**DOI:** 10.1101/2024.10.20.619279

**Authors:** Amy Luo

**Affiliations:** University of Tennessee, Knoxville

## Abstract

Ecologists and epidemiologists frequently rely on spatially distributed data. Studies in these fields may concern geographic boundaries, as environmental variation can determine the spatial distribution of organismal traits or diseases. In such cases, environmental boundaries produce coincident geographic boundaries in, for example, disease prevalence. Boundary analysis can be used to investigate the co-occurrence of organismal trait or disease boundaries and underlying environmental boundaries. Boundary and boundary overlap statistics test for the presence of significant geographic boundaries and spatial associations between the boundaries of two variables. There currently exists one implementation of boundary overlap statistics, though only on Windows and ESRI ArcView, limiting the availability of boundary overlap statistics to researchers. I have created BoundaryStats—an R package available on CRAN—that implements boundary and boundary overlap statistics. BoundaryStats is the first open-source, cross-platform implementation of these statistical methods making the statistics more widely accessible to researchers.

## 2 Introduction

Geographic boundaries are an intrinsic feature of spatial ecology and epidemiology, as the relationships between organismal traits or disease prevalence with an underlying environmental variable often produce coincident geographic boundaries. Boundaries are areas in which spatially distributed variables (e.g., bird plumage coloration, disease prevalence, annual rainfall) rapidly change over a narrow space. They can also represent edges or discontinuities (e.g., neighborhood edges, ecotype boundaries). Boundary zones themselves may be of interest; for example, the temporal boundary dynamics between ecotypes can provide insight into the factors that produce mosaic landscapes^1^.

Boundary analysis involves the assessment of whether significant geographic boundaries are present^2,3^ and whether the boundaries of multiple variables are spatially correlated^4^. Such analysis includes two categories of statistical test: boundary statistics (i.e., tests for the presence of cohesive boundaries) and boundary overlap statistics (i.e. tests for spatial association between boundaries). BoundaryStats runs two boundary statistics tests and three boundary overlap statistics, as initially described in Jacquez 1995^5^ and Fortin et al. 1996^6^. The boundary statistics are (1) the length of the longest boundary and (2) the number of cohesive boundaries on the landscape^6^. The boundary overlap statistics are (1) the amount of direct overlap between boundaries in variables A and B, (2) the mean minimum distance between boundaries in A and B, and (3) the mean minimum distance from boundaries in A to boundaries in B^5,6^.

While other spatial statistics account for complications like spatial autocorrelation and environmental heterogeneity^7^, boundary analysis can uniquely leverage geographic discontinuities to answer spatial questions. By identifying significant cohesive boundaries, researchers can delineate relevant geographic sampling units (e.g., populations as conservation units for a species or human communities with increased disease risk)^2^. Associations between the spatial boundaries of two variables can be useful in assessing the extent to which an underlying landscape variable drives the spatial distribution of a dependent variable. For example, ecologists are often interested in whether landscape-level ecological boundaries limit gene flow, thereby producing population structure; if the putative ecological boundary is limiting gene flow, one would expect concordant geographic boundaries in the landscape variable and population structure^3,8^. Landscape boundaries can similarly limit the distribution of taxonomically similar species^9^. In an epidemiological context, this may look like neighborhood effects on public health outcomes, including COVID-19 infection risk^10,11^ or spatial relationships between high pollutant density and increased disease risk^12,13^.

Currently, there is at least one tool that has implemented boundary overlap statistics: GEM, which was released as an extension of ESRI ArcView and a standalone Windows package^4^. GEM is not available as a cross-platform, free, and open-source software, thereby limiting its accessibility to researchers. R is a common, well-supported, and cross-platform language for statistical analysis. BoundaryStats implements boundary and boundary overlap statistics in R. It is available to download on the Comprehensive R Archive Network (CRAN), making the tools more accessible for researchers, especially in epidemiology and spatial ecology.

## 3 Boundary definitions

In this framework, we classify raster cells into a pseudobinary: boundary elements (1), non-boundary cells (0), or missing data (NA). For categorical variables, the algorithm for identifying boundary elements is simple: if any of a cell’s neighbors belongs to a different category, the cell classified as a boundary element. For quantitative variables, cells with the highest boundary intensity values, with a threshold set by the user, are classified as boundary elements. Boundaries are defined here as subgraphs of boundary elements, or contiguous cells that are all marked as boundary elements (Fig 1). For the purpose of defining subgraphs, cells are considered neighbors using the queen criterion (i.e., eight neighboring cells, including diagonally touching cells).

**Fig 1.**
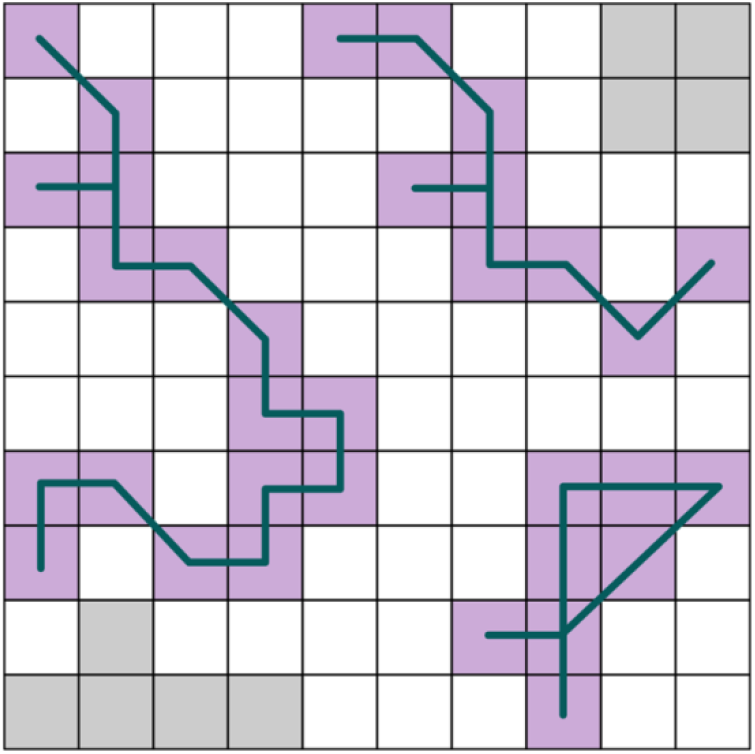
Example boundary subgraphs. Gray cells are missing values, white cells are non-boundary cells, and purple cells are boundary elements. Subgraphs are each represented using a line connecting all the boundary element cells that comprise them.

Boundary intensities for landscape-level variables can be calculated in a number of ways, including lattice- and triangulation-wombling^5,6,14,15^, fuzzy set modeling^5^, Monmonier’s algorithm^16^, spatial Bayesian clustering^17,18^, agglomeration of inner lines^19^, and removal of outer lines^19^. For quantitative variables, BoundaryStats will accept raster objects with the spatial variable directly or boundary values calculated from these or other methods. If given boundary intensity values, boundary elements will be classified directly using the top percent of values. The default proportion of values is 0.2, though this threshold can be changed by the user. When given the variable directly, BoundaryStats will use the Sobel-Feldman operator to calculate the boundary intensity. In accepting either the variables or boundary intensities, there is flexibility for users to define boundaries using relevant metrics for their data.

The Sobel-Feldman operator is commonly used for edge detection in computer vision applications. It approximates the magnitude of the partial derivative (i.e., rate of change) across each cell using the following kernels:

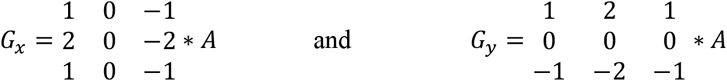

where *A* is the input raster cell, and G_x_ and G_y_ are the rates of variable change in the horizontal or vertical directions, respectively. The boundary intensity value is the overall rate of change:

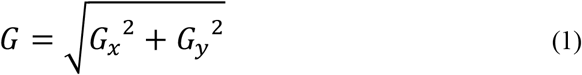

## 4 Statistics

BoundaryStats runs two boundary statistics and three boundary overlap statistics, as initially described in Jacquez 1995^5^ and Fortin et al. 1996^6^. Below, I describe these five statistics:

### Number of subgraphs

The first boundary statistic is the number of subgraphs. The number of subgraphs describes the number of boundaries on the landscape for a variable. In a raster of boundary elements, it is the number of unique subgraphs (three subgraphs each in fig 2a and fig 2b).

**Fig 2.**
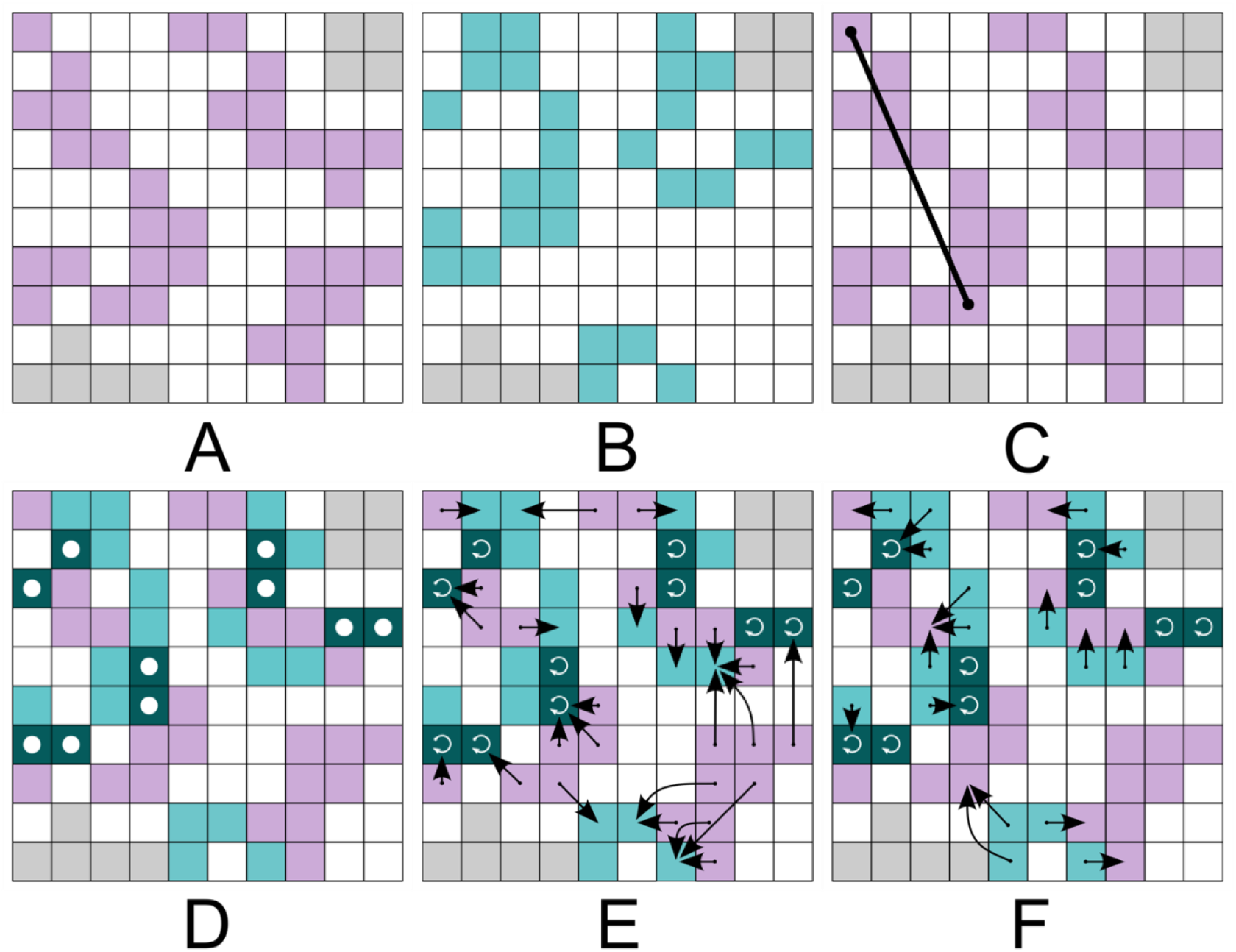
Example boundaries and statistics. (A) and (B) are boundary elements for hypothetical variables A and B. White cells are non-boundaries, gray are missing values, and purple or teal are boundary elements. (C) Length of the longest subgraph. (D) Produced by overlapping cells in A and B. Dark blue cells, highlighted by white dots, are where the boundary elements overlap. (E) For every boundary element for variable A, the nearest boundary element for variable B. Circular arrows indicate distance to self. (F) For every boundary element for B, the nearest boundary element for A.

### Longest subgraph

The other boundary statistic included here is the length (in meters) of the longest subgraph, or boundary. The function calculates the longest length across each subgraph, then converts the length to distance based on the cell resolution and the projection of the raster. The length of the longest subgraph is then retained (Fig 2c).

### Direct overlap

The direct overlap statistic, *O*_*d*_, is a count of the number of overlapping boundary elements of two variables (Fig 2d).

### Mean minimum distance between boundaries

This statistic describes the spatial proximity between boundaries of variables x and y, as defined by the mean distance to the nearest boundary element of the other variable. Spatial relationships between boundaries may not result in direct overlap, so this statistic accounts for potential correlations in non-overlapping boundaries. For each boundary element in variable x, the function calculates the distance to the nearest boundary element in variable y, then repeats the inverse for each boundary element in variable y (Fig 2e and 2f). It then takes the mean of these minimum distances across boundary elements in x and y:

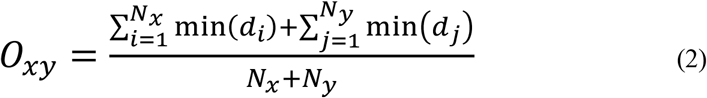

where *i* and *j* are boundary elements for variables *x* and *y*, respectively; *min(d)* is the minimum distance between a boundary element for one variable to a boundary element in the other variable; and *N* is the number of boundary elements for the variable.

### Mean minimum distance from boundary x to boundary y

This statistic describes the mean distance from boundary elements in x to the nearest boundary element in y. It is an indicator for whether the boundaries in variable x depend on variable y. The reciprocal nature of the previous statistic implies some reciprocity of effect, as opposed to the unidirectionality implicit here. For each boundary element in the raster for x, the function calculates the distance to the nearest boundary element of variable y, then takes the mean across all boundary elements in x (Fig 2e):

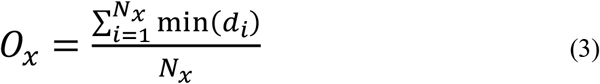

## 5 Neutral models

In addition to calculating each statistic, BoundaryStats uses iterations of a neutral landscape model to determine whether the boundaries in the input landscape differ from a random landscape. Users select a neutral landscape model and number of iterations of that model to produce a null distribution of each statistic, based on the selected model and the structure of the input landscape. BoundaryStats implements three neutral landscape models: stochastic landscapes, Gaussian random fields, and modified random clusters.

The simplest neutral landscape model is complete stochasticity. It takes all the cell values from the input raster and assigns each value to a random cell. Each cell in the simulated raster is assigned a value from the original dataset, with no replacement of values. It will ignore cells with missing data (e.g., for terrestrial data along a coastline, it will not draw from or assign new values to ocean cells). The simulated raster has the exact same values as the original raster, but values are randomly placed with no spatial autocorrelation.

The next neutral landscape model simulates a Gaussian random field with the same degree of spatial autocorrelation as the input raster. It is suited for continuous or discrete quantitative variables. This method calculates the local Moran’s I across the original raster and builds local indication of spatial association (LISA) clusters^20^. Each cluster circumscribes an area with significant local spatial autocorrelation. The maximum distances across each LISA cluster are extracted, then the median is taken across clusters; the resulting value represents the average range of significant spatial autocorrelation. The function then simulates a Gaussian random field with the same range of spatial autocorrelation, extent, and resolution as the input raster. If the spatial autocorrelation range is too large for the spatial extent, this parameter is reduced by 10% and another Gaussian random field is attempted; this step is repeated until a simulated raster is successfully produced. If there are missing data cells in the original landscape (e.g., a coastline or lake is present, but the data are terrestrial), the corresponding cells in the simulated raster will be set as missing data (NA).

The modified random cluster model is an implementation of the method described by Saura and Martínez-Millan for simulating neutral landscapes for categorical variables^21^. The first step is a percolated raster (Fig 3a). Each cell is assigned a value 0 ≤ *x* ≤ 1 from a uniform distribution, and cells with values above a threshold probability *p* are marked. *p* is defined by the user, and higher values of *p* result in larger cluster sizes in the final simulated raster. Next, contiguous sets of marked cells are grouped into clusters, using the rook criterion (i.e., neighbors are the four edge-touching neighbors) (Fig 3b). Clusters are then assigned a category (Fig 3c). Categories from the input raster are chosen one at a time, and random clusters are assigned to that category. When the proportion of that category in the simulated raster reaches the proportion in the original raster, clusters are then assigned to the next category, until all the clusters are assigned. In the last step, the unmarked cells are categorized based on the most frequent category among their neighbors using the queen criterion (Fig 3d). If there is a tie between two categories, one of the tied categories is picked at random. If all neighbors are unassigned, a random category is picked; probabilities for each category are based on their proportions in the input raster. The simulated raster is then cropped, so that the simulated raster has missing data in the same areas as the input raster.

**Fig 3.**
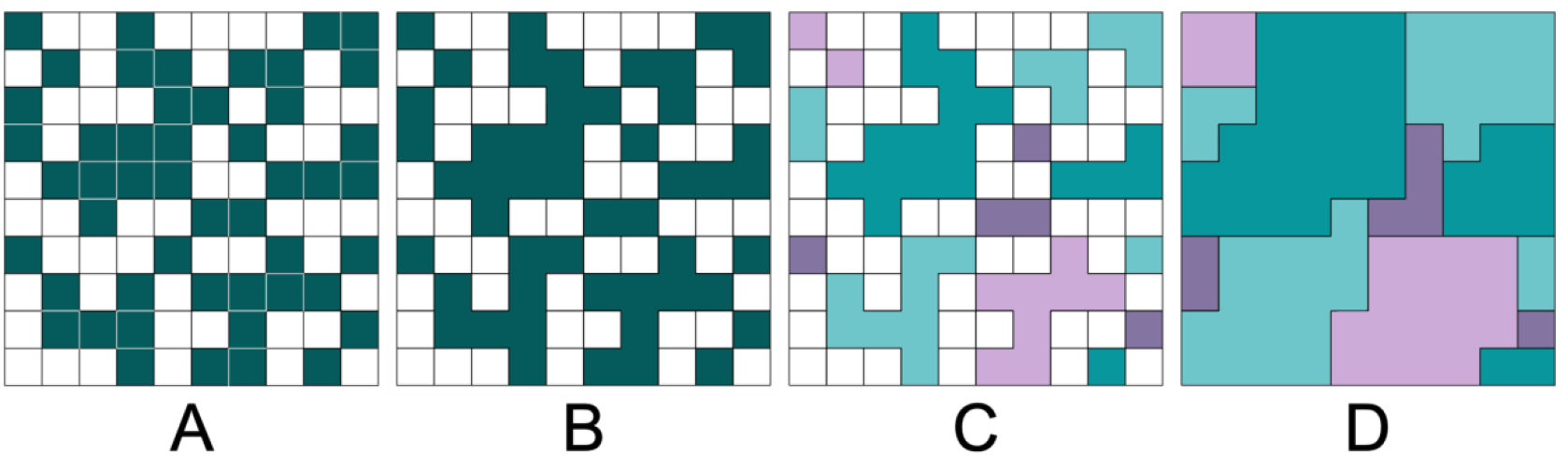
Modified random cluster procedure, adapted from Saura and Martínez-Millan 2000. (A) Percolated raster with *p* = 0.5. (B) Marked cells merged into clusters. (C) Clusters assigned a category. (D) Unmarked cells filled based on neighbors.

## 6 Implementation and Example

Data in this example are from Luo et al.^22^, in which the authors hypothesized that song divergence is facilitating genetic divergence in white-crowned sparrows through speciation by sexual selection. The data below are song boundaries and genetic admixture interpolations from the study. The song data are boundary intensity values estimated using GeoOrigins^23^, based on the acoustic dissimilarity and spatial relationship between recorded songs. The genetic data are spatially interpolated admixture coefficients. The admixture coefficients of samples were estimated in fastSTRUCTURE^24^ and interpolated using local kriging.

### Read in data

Read in raster data to terra^25^ SpatRaster objects (Figure 4). The two objects need the same projection, extent, and resolution.

~~~
>library(terra)
>library(magrittr)
>songs <-rast(‘2010_2022_song_boundaries.asc’)
>genetic <-rast(‘genetic_interpolation.asc’) %>%
resample(., songs)
>songs <-crop(songs, genetic) %>%
mask(., genetic)
~~~

**Fig 4.**
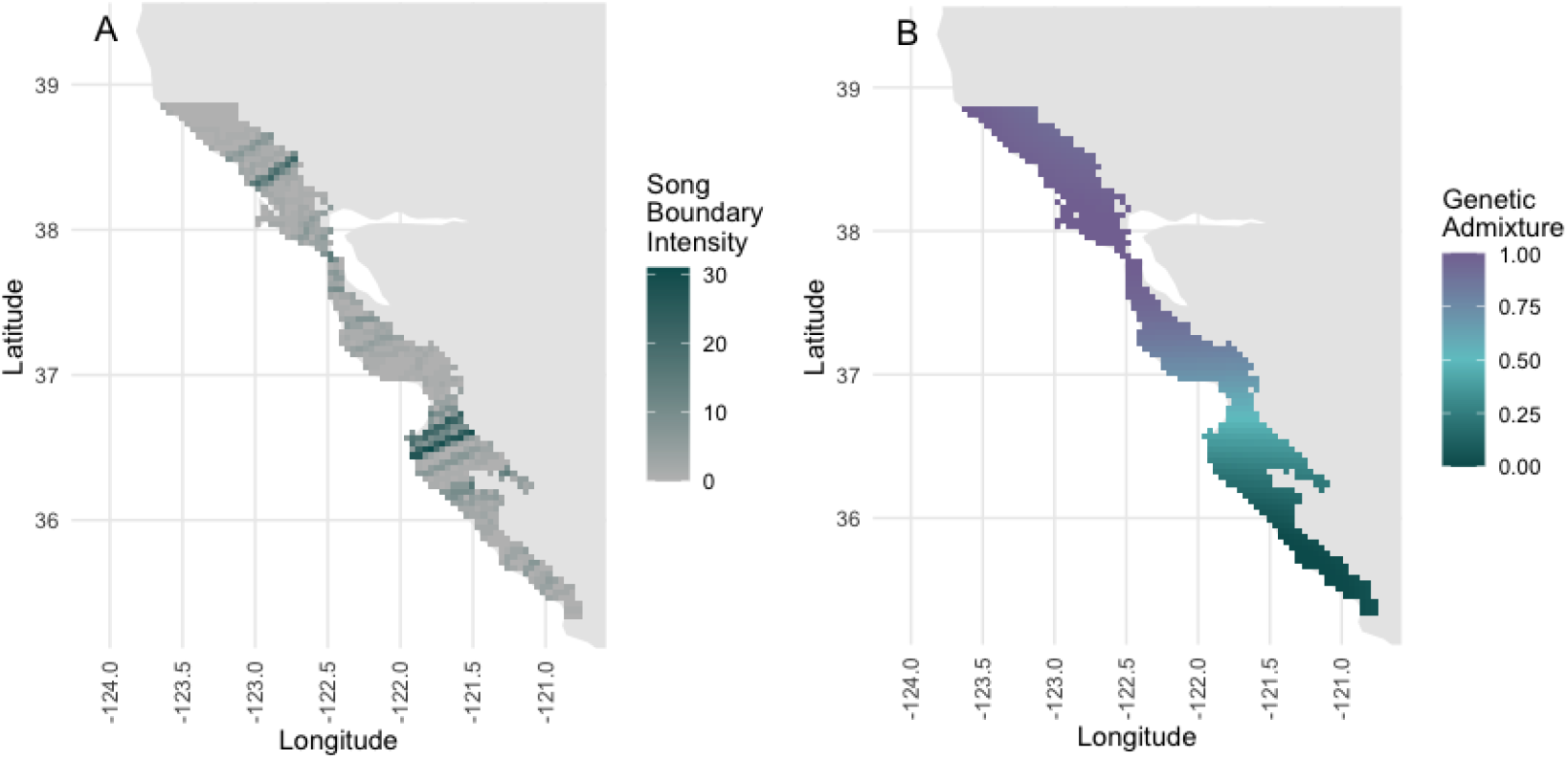
Maps of (A) song boundary intensity and (B) genetic admixture between two populations.

### Calculate spatial boundaries for variables

If the variable is categorical, use the *categorical_boundary* function. For continuous variables, use the *define_boundary*, which applies a proportion threshold for the highest boundary intensity values (default = 0.2). Boundary intensity can be calculated however the user chooses; if the input raster for a continuous variable already contains boundary intensities, the argument *convert* in *define_boundaries* should be set to FALSE (default). For a raster with the variable, one can set *convert* to TRUE to use the Sobel-Feldman operator to calculate boundary intensities.

Both variables in this example are continuous, so I will use *define_boundary*. The song raster already contains boundary intensity values, from which boundary elements can be directly determined, so I can use the default of FALSE for *convert*. But the genetic data are spatially interpolated from the admixture coefficients of sampled white-crowned sparrows, so boundary intensity needs to be calculated from this variable (*convert* = TRUE).

~~~
>library(BoundaryStats)
>song_boundaries <-define_boundary(songs)
>genetic_boundaries <-define_boundary(genetic, convert = TRUE)
~~~

### Plot boundary overlap

This optional step is to visualize where the boundaries of the two variables are overlapping using *plot_boundary* (Figure 5). The function is a wrapper function for ggplot^26^, and the colors and trait names can optionally be customized.

**Fig 5.**
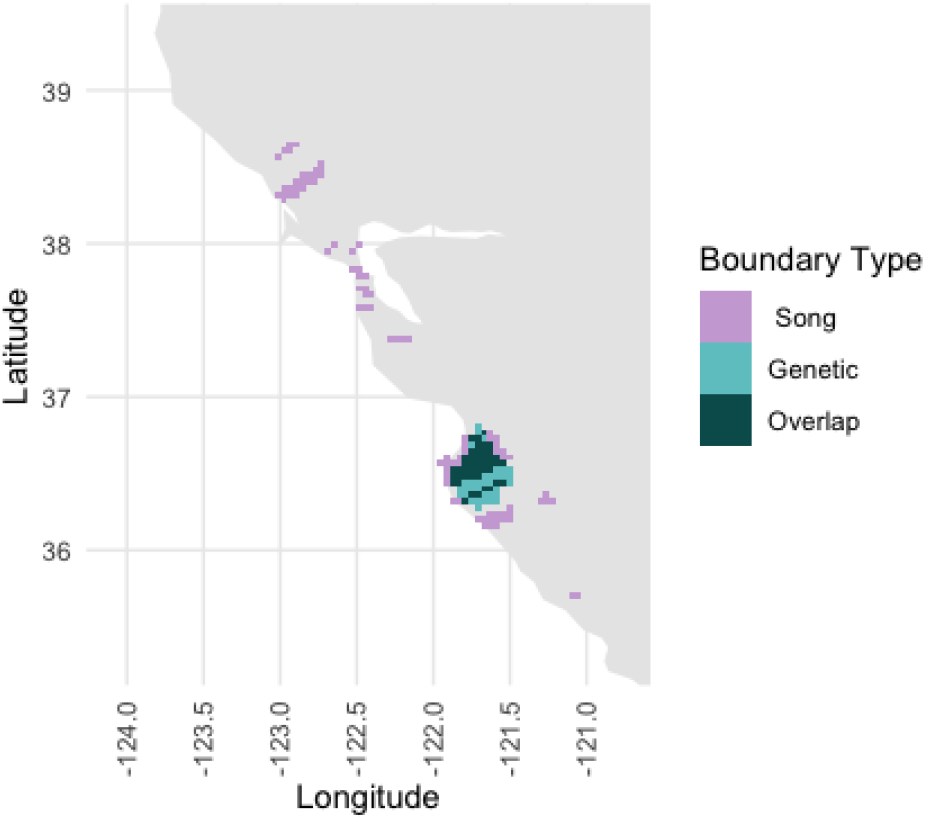
Output of the plot_boundary function.

### Create null distributions for statistics

For both boundary statistics, use *boundary_null_distrib*. For the three overlap statistics, use *overlap_null_distrib*. Both functions simulate random iterations of a raster based on the specified neutral landscape model and input data. Statistics are calculated for each iteration. Custom null probability distributions are calculated based on the iterations.

The functions take the SpatRaster object(s) containing the landscape variable, a neutral landscape model, whether the variable is categorical, and the number of iterations. For *overlap_null_distrib*, separate models can be specified for the two variables, and the first raster is assumed to depend on the second raster. The argument *rand_both* specifies whether y should be modeled in each iteration; since some hypotheses assume variable x depends on an underlying distribution of boundaries in variable y, users can choose to keep boundaries in y static for each iteration. For this example, the genetic boundary is hypothesized to depend on song boundaries. Therefore, the SpatRaster object containing the genetic admixture interpolation is the first argument, and I keep variable y static (*rand_both* = FALSE).

~~~
>song_boundary_null <-boundary_null_distrib(songs, convert = FALSE, cat = FALSE,
n_iterations = 1000, threshold = 0.2, ‘gaussian’)
>genetic_boundary_null <-boundary_null_distrib(genetic, convert = TRUE, cat = FALSE,
n_iterations = 1000, threshold = 0.2, ‘gaussian’)
>boundary_overlap_null <-overlap_null_distrib(genetic, songs, rand_both = FALSE,
n_iterations = 1000, x_convert = TRUE, threshold = 0.2, x_model = ‘gaussian’)
~~~

### Run statistical tests

Both boundary statistics functions require only the raster with boundary elements for the variable and the matching null distribution object, produced by *boundary_null_distrib*

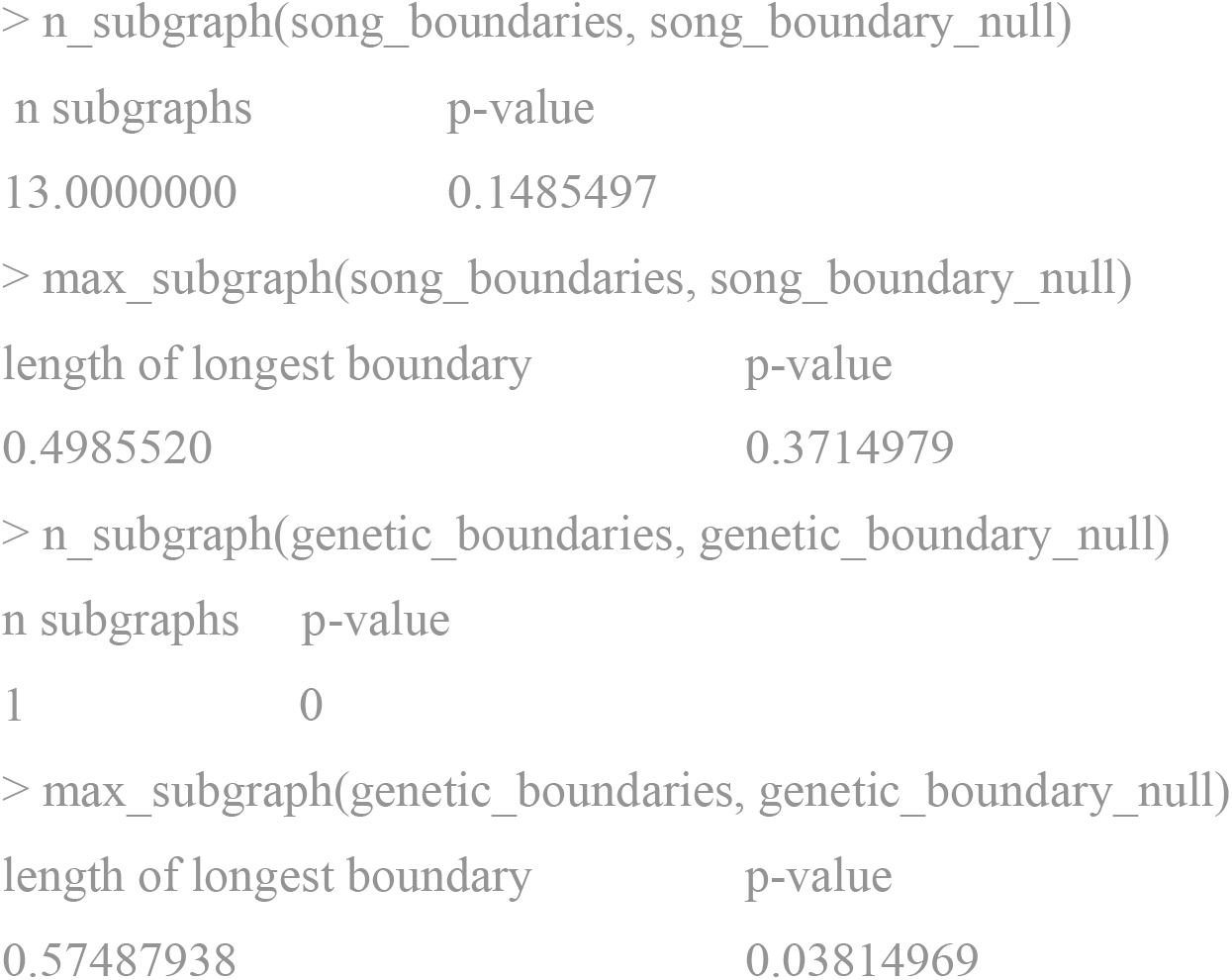

The boundary overlap statistics similarly takes the boundary element raster and null distribution as arguments. In this case, it requires two boundary element SpatRaster objects, one for each variable. The order of the variables should match that used in *overlap_null_distrib*. In this case, genetic boundaries first, then song boundaries.

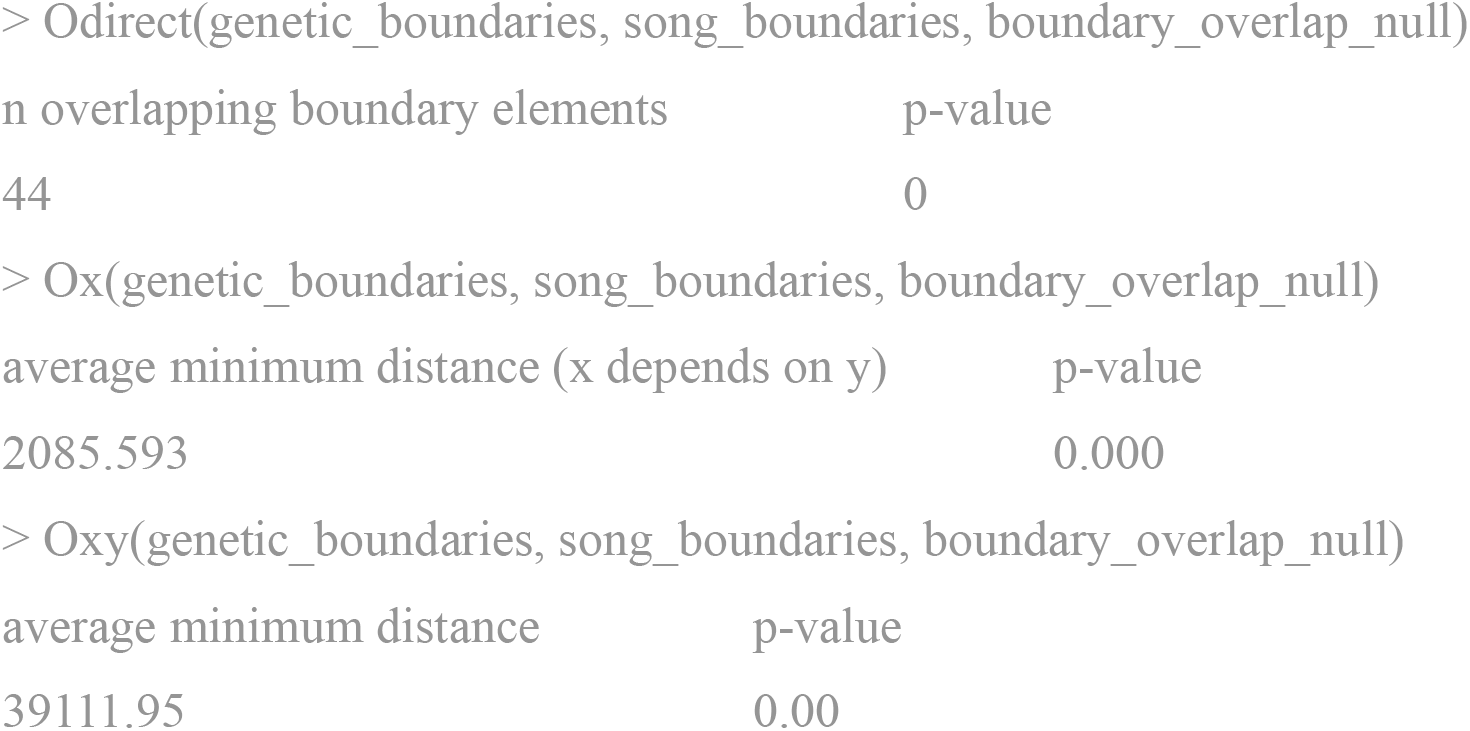

### Interpretation of example data output

When analyzing the data from Luo et al.^22^, all the statistical tests recovered a significant result, most with p-values so small they were rounded down to 0. The results suggest the presence of cohesive genetic boundary and song boundaries, rather than artifacts of spatial noise. Further, the genetic boundary has significant direct overlap and spatial proximity to a song boundary. While boundary overlap statistics can only demonstrate a correlation between boundaries, the results support the hypothesis that song boundaries are facilitating a coincident genetic boundary.

## 7 Summary

BoundaryStats implements five boundary and boundary overlap statistics that can be used for boundary analysis. Boundary analyses can be used across many contexts that make use of spatially distributed data. For example, spatial ecologists and epidemiologists can use boundary overlap statistics to assess whether environmental variables are influencing the distribution of organismal traits or disease occurrences. Environmental influences can, in some cases, be detected through the co-occurrence and coincidence of geographic boundaries; environmental boundaries may produce boundaries in the variables of interest. As such, this new open-source, cross-platform implementation will make boundary statistical methods more widely accessible to researchers.

